# Heterosis indices: What do they really measure?

**DOI:** 10.1101/800441

**Authors:** Dominique de Vienne, Julie B. Fiévet

## Abstract

Heterosis (hybrid vigour) is a universal phenomenon of crucial agro-economic and evolutionary importance. We show that the most common heterosis indices do not properly measure deviation from additivity because they include both a component accounting for “real” heterosis and a term that is not related to heterosis since it is derived solely from parental values. Therefore, these indices are inadequate when the aim of the study is to compare heterosis levels between different traits, environments, genetic backgrounds or developmental stages, as these factors may affect not only heterosis but also parental values. The only relevant index for such comparisons is the so-called “potence ratio”. These observations argue for the careful choice of heterosis indices depending on the purpose of the work.

**Highlight:** Unlike dominance indices, heterosis indices, with one exception, do not properly measure the level of deviation from additivity, thus making them unsuitable for comparative analyses.

## Introduction

Non-linear processes are extremely common in biology. In particular, genotype-phenotype or phenotype-phenotype relationships frequently display concave behaviours, resulting in the dominance of “high” over “low” alleles (Wright, 1934) and in positive heterosis for a wide range of polygenic traits (Fievet *et al*., 2018; Vasseur *et al*., 2019). Properly quantifying the degree of non-additivity is an essential prerequisite for interpreting and comparing genetic studies and for making predictions in plant and animal breeding. However, most of the classical heterosis indices do not meet this requirement.

First, let us recap how the degree of dominance is measured. There are two classical dominance indices:

(i) Wright (1934) defined:

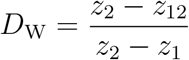

where *z*_1_, *z*_2_ and *z*_12_ are respectively the phenotypic values of genotypes *A*_1_*A*_1_, *A*_2_*A*_2_ and *A*_1_*A*_2_, with *z*_2_ > *z*_1_. *D*_W_ varies from 0, when *A*_2_ is fully dominant over *A*_1_, to 1, when *A*_2_ is fully recessive with respect to *A*_1_. *D*_W_ = 0.5 corresponds to semi-dominance or additivity 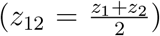 (Table 1). Note that D_W_ is strictly equivalent to the dominance coefficient h used in evolutionary genetics (Crow & Kimura, 1970).

(ii) Falconer (1960) proposed the following index:

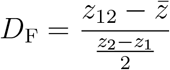

where 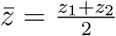. *D*_F_ varies in the opposite direction to *D*_W_: its value is 1 if *z*_12_ = *z*_2_ (complete dominance of *A*_2_ over *A*_1_), –1 if *z*_12_ = *z*_1_ (*A*_2_ is fully recessive with respect to *A*_1_) and 0 if there is additivity. In the case of overdominance, *D*_W_ < 0 and *D*_F_ > 1, and in the case of underdominance, *D*_W_ > 1 and *D*_F_ < –1 (Table 1).

**Table 1.**
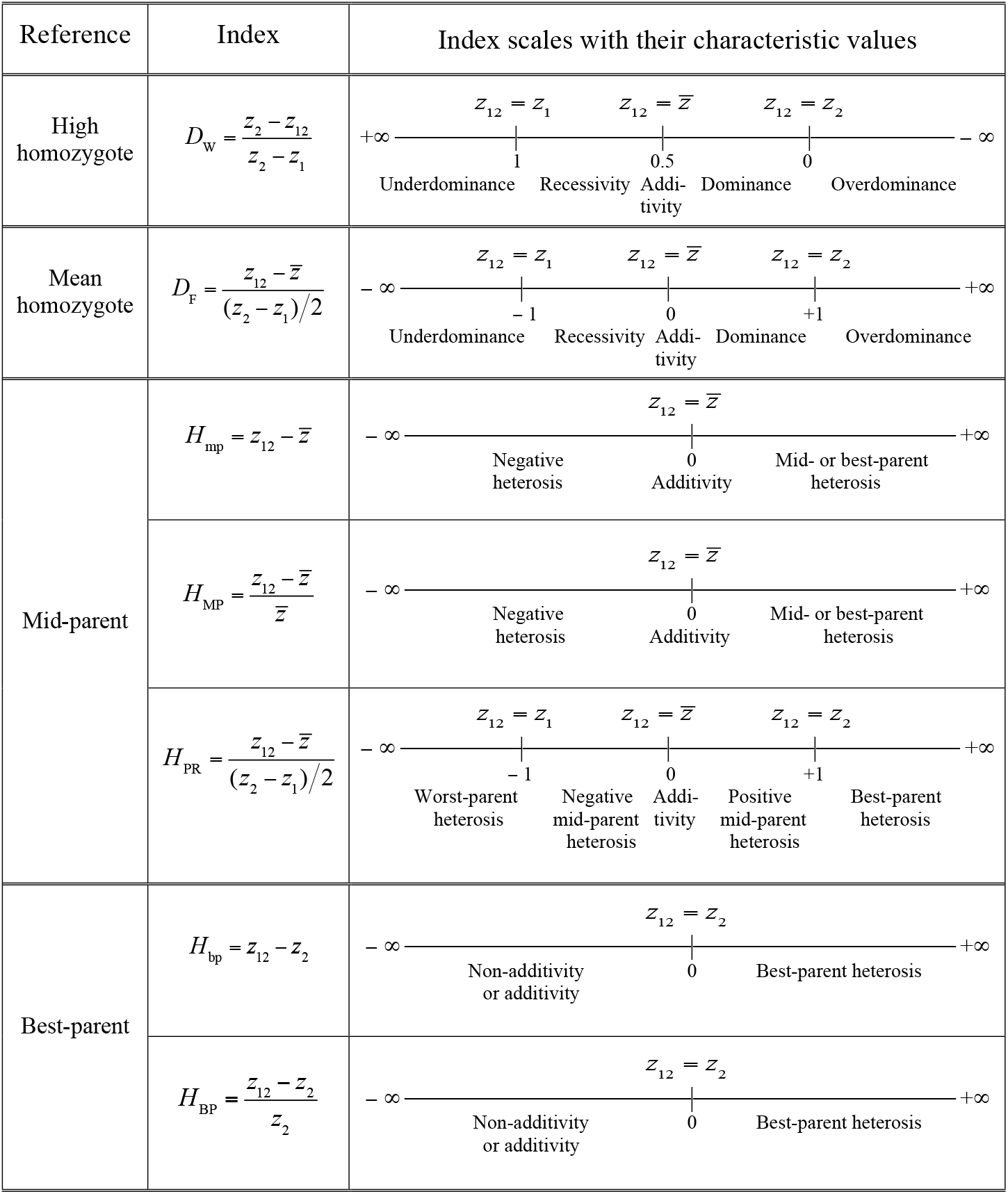
Dominance and heterosis indices. *D*_W_: Wright’s dominance index. *D*_F_: Falconer’s dominance index. *H*_mp_, *H*_MP_, *H*_PR_, *H*_bp_ and *H*_BP_: heterosis indices. Subscripts: mp or MP, mid-parent; PR, potence ratio; bp or BP, best-parent. In the equations of the text, *H*_PR_ is also noted *h*_P_ for simplicity. *z*_1_ (resp. *z*_2_): the phenotypic value of parental homozygote 1 or of parent 1 (resp. 2). *z*_12_: the heterozygote or hybrid value. 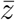: the mean parental value. By convention, *z*_2_ > *z*_1_.

| Reference | Index | Index scales with their characteristic values |
| --- | --- | --- |
| High homozygote | $D_W = \frac{z_2 - z_{12}}{z_2 - z_1}$ | $+\infty \text{ --- } \frac{z_{12} = z_1}{1} \text{ --- } \frac{z_{12} = \bar{z}}{0.5} \text{ --- } \frac{z_{12} = z_2}{0} \text{ --- } -\infty$<br>Underdominance Recessivity Additivity Dominance Overdominance |
| Mean homozygote | $D_F = \frac{z_{12} - \bar{z}}{(z_2 - z_1)/2}$ | $-\infty \text{ --- } \frac{z_{12} = z_1}{-1} \text{ --- } \frac{z_{12} = \bar{z}}{0} \text{ --- } \frac{z_{12} = z_2}{+1} \text{ --- } +\infty$<br>Underdominance Recessivity Additivity Dominance Overdominance |
| Mid-parent | $H_{mp} = z_{12} - \bar{z}$ | $-\infty \text{ --- } \frac{z_{12} = \bar{z}}{0} \text{ --- } +\infty$<br>Negative heterosis Additivity Mid- or best-parent heterosis |
| | $H_{MP} = \frac{z_{12} - \bar{z}}{\bar{z}}$ | $-\infty \text{ --- } \frac{z_{12} = \bar{z}}{0} \text{ --- } +\infty$<br>Negative heterosis Additivity Mid- or best-parent heterosis |
| | $H_{PR} = \frac{z_{12} - \bar{z}}{(z_2 - z_1)/2}$ | $-\infty \text{ --- } \frac{z_{12} = z_1}{-1} \text{ --- } \frac{z_{12} = \bar{z}}{0} \text{ --- } \frac{z_{12} = z_2}{+1} \text{ --- } +\infty$<br>Worst-parent heterosis Negative mid-parent heterosis Additivity Positive mid-parent heterosis Best-parent heterosis |
| Best-parent | $H_{bp} = z_{12} - z_2$ | $-\infty \text{ --- } \frac{z_{12} = z_2}{0} \text{ --- } +\infty$<br>Non-additivity or additivity Best-parent heterosis |
| | $H_{BP} = \frac{z_{12} - z_2}{z_2}$ | $-\infty \text{ --- } \frac{z_{12} = z_2}{0} \text{ --- } +\infty$<br>Non-additivity or additivity Best-parent heterosis |

The *D*_W_ and *D*_F_ indices are linearly related:

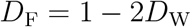

Thus, dominance can be quantified with either index, both of them giving the position of the heterozygote relative to the parental homozygotes. The two indices merely vary in opposite directions, with *D*_W_ decreasing when dominance increases.

For polygenic traits, either index could be used to quantify non-additivity, *i.e*. real heterosis, without any ambiguity. Actually, one finds five heterosis indices in the literature (see their characteristic values in Table 1).

The two most popular indices are the best-parent (BP) and mid-parent (MP) heterosis indices (e.g. Gowen, 1952; Frankel, 1983):

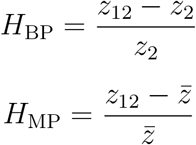

where *z*_2_, *z_12_* and 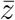 are respectively the phenotypic values of the parent 2 (with *z*_2_ > *z*_1_), of the parent 1 × parent 2 hybrid and of the parental mean.

In some instances, the authors do not normalize the difference between the hybrid and the best-or mid-parent value:

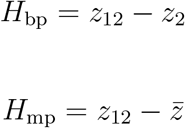

Finally, the so-called “potence ratio” (Mather, 1949) has the same expression as Falconer’s dominance index:

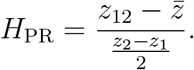

A value of 0 indicates additivity, 1 indicates *z*_12_ = *z*_2_ (hybrid value = best-parent value), –1 indicates *z*_12_ = *z*_i_ (hybrid value = worst-parent value) and > 1 (resp. < –1) indicates best-parent (resp. worst-parent) heterosis (Table 1). *H*_PR_ includes the values of the three genotypes, whereas the other indices lack one of the parental values (*H*_BP_ and *H*_bp_) or both (*H*_MP_ and *H*_mp_). From a genetic point of view, *H*_PR_ is explicitly expressed in terms of the five genetic effects contributing to heterosis (Supplementary Table S1). Thus, the potence ratio, which is still by far the least used heterosis index, is the only one that informs us of the exact position of the hybrid value relative to the parental values. Wright’s dominance index has the same property, but its inverse direction of variation, which makes comparisons less easy, probably explains why it is not used in this context.

## Relationship between the potence ratio and the other heterosis indices

It is easy to show that the relationship between *H*_PR_, hereafter noted *h_P_* for simplicity, and the other indices is (with *z*_2_ > *z*_i_):

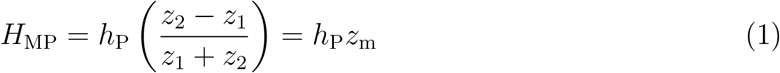

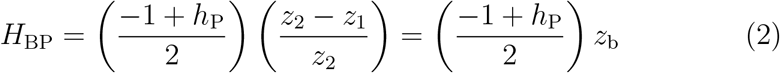

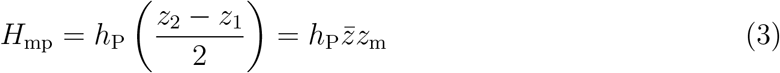

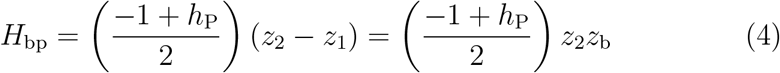

where 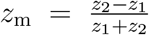 is the variation coefficient (*σ*/*μ*) of the trait in the parents and 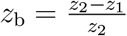 is the difference between the parents normalized by the best-parent value (or *z*_b_ = 2*σ*/*z*_2_).

For a given *h*_P_ value, the indices *H*_MP_ and *H*_BP_ are linearly related to *z*_m_ and *z*_b_, respectively, i.e. they depend on the scale of the parental values. More specifically, the relationship between *H*_MP_ and *z*_m_ is negative when *h*_P_ < 0 and positive when *h*_P_ > 0, while the relationship between *H*_BP_ and *z*_b_ is negative when *h*_P_ < 1 and positive when *h*_P_ > 1. As *z*_m_ and *z*_b_ are positive, we see from equation 1 that for *h*_P_ ≠ 0, we have

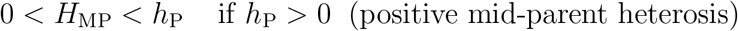

and

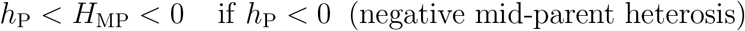

Regarding *H*_BP_, we see from equation 2 that for *h*_P_ ≠ 1, we have

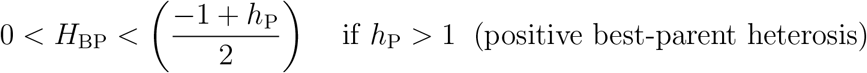

and

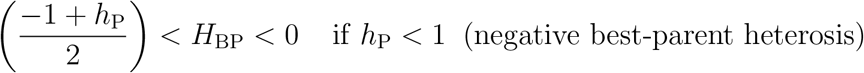

If *h*_P_ = 0 (resp. *h*_P_ = 1), *H*_MP_ (resp. *H*_BP_) = 0.

Numerical applications performed with nine *h*_P_ values, from *h*_P_ = –2 to *h*_P_ = 2, show that the same *H*_MP_ or *H_BP_* value can be observed for very different *h*_P_ values (Supplementary Figure S1). For instance, *H*_MP_ ≈ 0.4 can correspond to both mid-parent heterosis (*h*_P_ = 0.5, *z*_m_ ≈ 0.8) and best-parent heterosis (*h*_P_ = 2, *z*_m_ ≈ 0.21).

We illustrate this using experimental data from maize. We measured six traits (flowering time, plant height, ear height, grain yield, thousand-kernel weight and kernel moisture) in four crosses (B73×F252, F2×EP1, F252×EP1, F2×F252) grown in three different environments in France (Saint-Martin-de-Hinx in 2014, Jargeau in 2015 and Rhodon in 2015). We computed *h*_P_, *H*_MP_ and *H*_BP_ for the 72 trait-cross-environment combinations. Fig. 1A,B shows that the relationship between *h*_P_ and the other two indices is very loose, if any. A given *h*_P_ value can correspond to a wide range of *H*_MP_ or *H*_BP_ values, and vice versa. We performed the same analyses using the data published by Shang *et al*. (2016), who measured five traits in two crosses of cotton grown in three environments. The same loose relationship between *h*_P_ and either heterosis index was observed (Fig. 1C,D). This means that the trait variation coefficients or the normalized difference between the parents, which are not related to heterosis since they do not include values from the hybrids, markedly affect *H*_MP_ and *H*_BP_.

**Figure 1.**
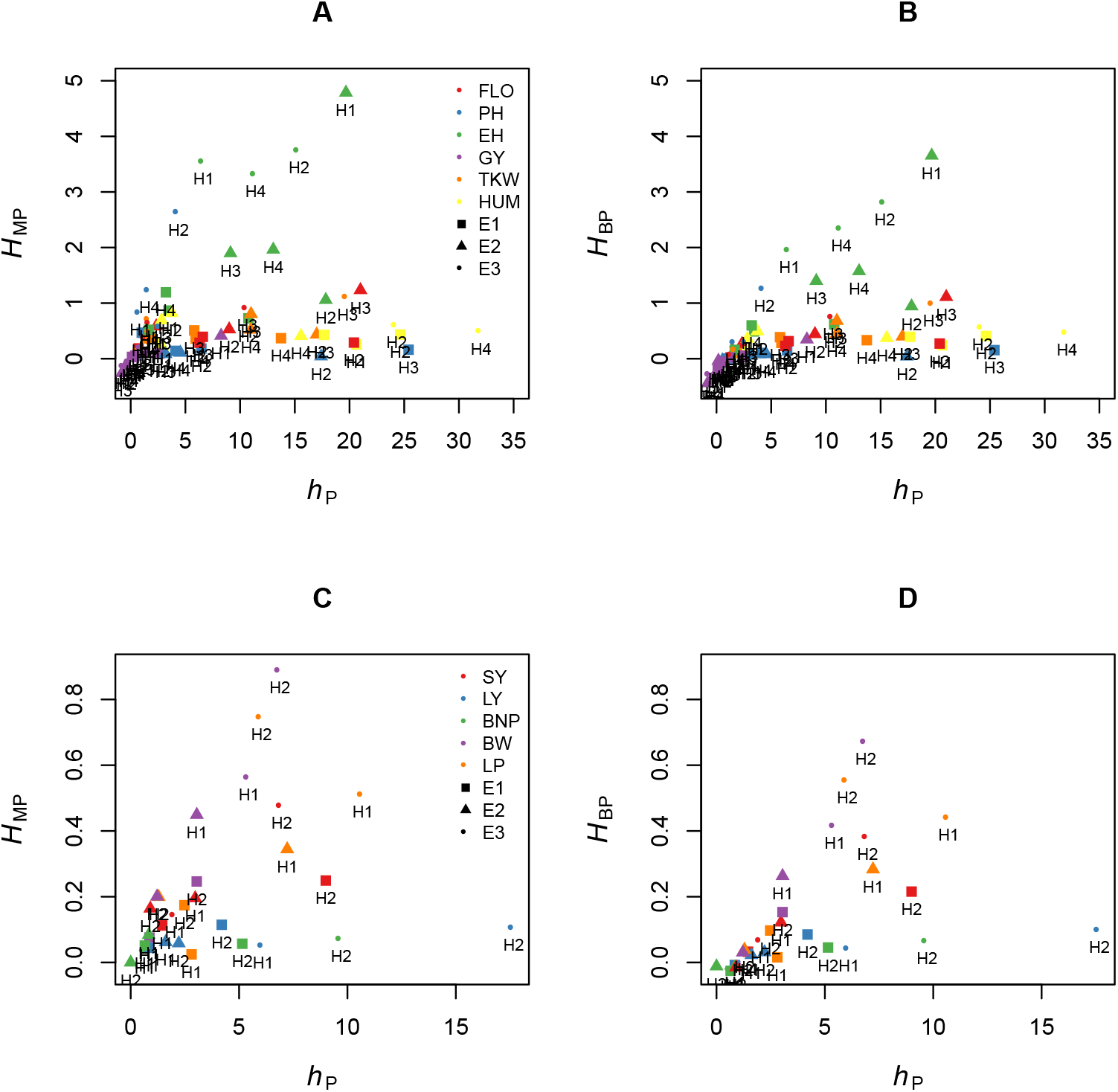
Relationship between the potence ratio *h*_P_ and the two heterosis indices *H*_MP_ and *H*_BP_. (A) & (B) Six traits were measured in maize (FLO: flowering time [days between 50 % flowering and August, 12^th^], PH: plant height, EH: ear height, GY: grain yield, TKW: thousand-kernel weight, KM: kernel moisture) in four crosses (H1: B73×F252, H2: F2×EP1, H3: F252×EP1, H4: F2×F252) grown in three environments in France (E1: Saint-Martin-de-Hinx, E2: Jargeau, E3: Rhodon). (A) Relationship between *h*_P_ and *H*_MP_. (B) Relationship between *h*_P_ and *H*_BP_. (For clarity, four of the 72 trait-cross-environment combinations are not represented because they have high hP values.) (C) & (D) Five traits were measured in cotton (SY: seed yield, LY: lint yield, BNP: bolls per plant, BW: boll weight, LP: lint percent) in two crosses (H1: X1135×GX100-2 and H2: GX1135×VGX100-2) grown in three environments in China (E1: Handan, E2: Cangzhou, E3: Xiangyang) (Data from Shang *et al*., 2016). (C) Relationship between *h*_P_ and *H*_MP_. (D) Relationship between *h*_P_ and Hbp.

Regarding the *H*_MP_ and *H*_BP_ indices, which are not dimensionless, they only provide the direction of heterosis. For a given *h*_P_ value, *H*_MP_ can vary from –∞ to 0 when *h*_P_ < 0 and from 0 to +∞ when *h*_P_ > 0, and *H*_BP_ can vary from – ∞ to 0 when *h*_P_ < 1 and from 0 to +∞ when *h*_P_ > 1 (equations 3 and 4).

Let us examine the possible interpretation errors that may result from the use of the most common heterosis indices.

## The pitfalls of the most commonly used heterosis indices

The non-univocal relationship between *h*_P_ and the most commonly used heterosis indices has two consequences. (i) Comparing index values for a given trait in different crosses and/or environments and/or developmental stages leads to erroneous conclusions whenever these factors have an effect on the scale of the trait and/or on the difference between the parental values (*i.e*. on *z*_m_ or *z*_b_). Possible differences in actual heterosis levels between these conditions cannot be detected. (ii) This problem is even more pronounced when studying different traits because each trait has its own scale of variation, making *H*_MP_ and *H*_BP_ (and to a greater extent *H*_mp_ and H_bp_) useless for comparing the real levels of heterosis of these traits.

These pitfalls can easily be illustrated from our maize dataset. Fig. 2A shows that classifying a set of traits according to their degree of heterosis can give markedly different results depending on whether one uses the *h*_P_ index or one of the two indices *H*_MP_ and H_BP_. For instance, in the F252×EP1 cross, flowering time displays moderate heterosis according to *H*_MP_ and *H*_BP_ even though this trait actually has the highest *h*_P_ value. Conversely, plant height is the second most heterotic trait according to *H*_MP_ and *H*_BP_, but not according to *h*_P_. Similarly, comparing heterosis of a given trait in different hybrids results in index-specific rankings: heterosis of ear height measured with *h*_P_ is highest in the B73×F252 hybrid, whereas according to *H*_MP_ and *H*_BP_ the highest values are found in the F252×EP1 hybrid (Fig. 2B). Finally, the effect of the environment on heterosis reveals the same discrepancies between *h*_P_ on the one hand and *H*_MP_ or *H*_BP_ on the other hand (Fig. 2C).

**Figure 2.**
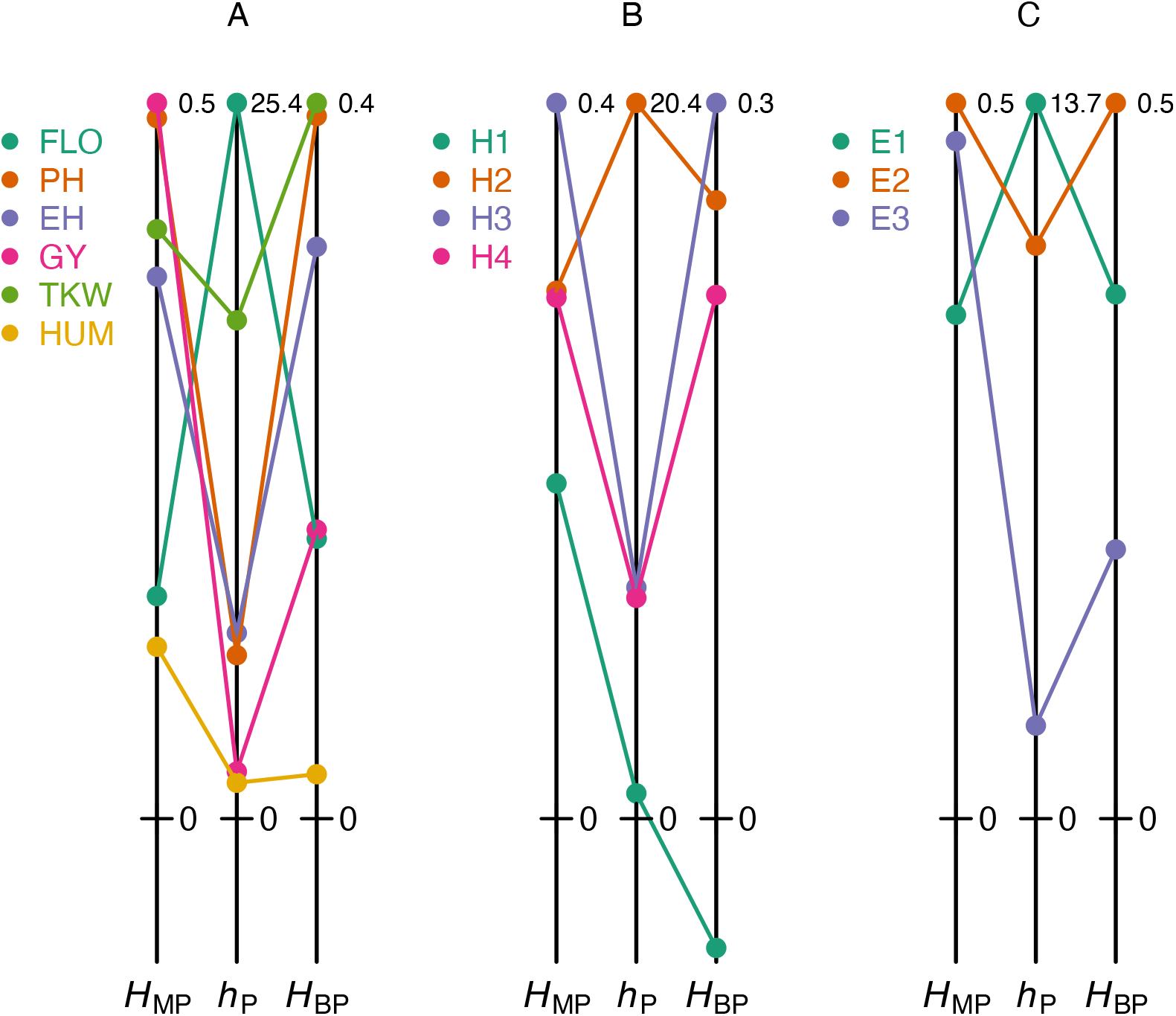
Heterosis values calculated using different indices. (A) Heterosis indices for six traits measured in the F252×EP1 cross grown in Saint-Martin-de-Hinx (France) in 2014. (B) Heterosis indices for ear height in four crosses grown in Saint-Martin-de-Hinx (France) in 2014. (C) Heterosis indices for plant height in the F2×F252 cross grown in the three environments. The six traits and the three environments are the same as in Fig. 1A. The scales of the heterosis indices are normalized by the maximum value in each dataset (figures at the top right of the vertical lines).

It is also informative to compare the variation of heterosis indices for a trait measured during development or growth. We fitted the percentage of flowering individuals over time in the W117×F192 and W117×F252 hybrids and their parents, using the Hill function:

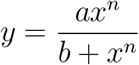

where *n* is the Hill coefficient. We then computed the variation of heterosis for the percentage of flowering individuals estimated from the fitted curves (Fig. 3). Again, *h*_P_ tells a different story compared to *H*_MP_ and *H*_BP_. Both *H*_MP_ and *H*_BP_ decrease as flowering progresses because the variation coefficient also decreases. This masks the evolution of real heterosis, which actually increases.

Similar results were observed in a simulation describing the increase in population size of a unicellular organism, which exhibits logistic growth.

We used:

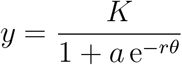

where *y* is the size of the population, *K* the carrying capacity, a a constant, *r* the growth rate and *θ* the time. We assumed that the parents differed only in their growth rate *r* and that there is additivity for this parameter. Results show that the variations over time of *H*_MP_ and *H*_BP_ index values for population size are clearly not congruent with that of *h*_P_ (Supplementary Figure S2).

**Figure 3.**
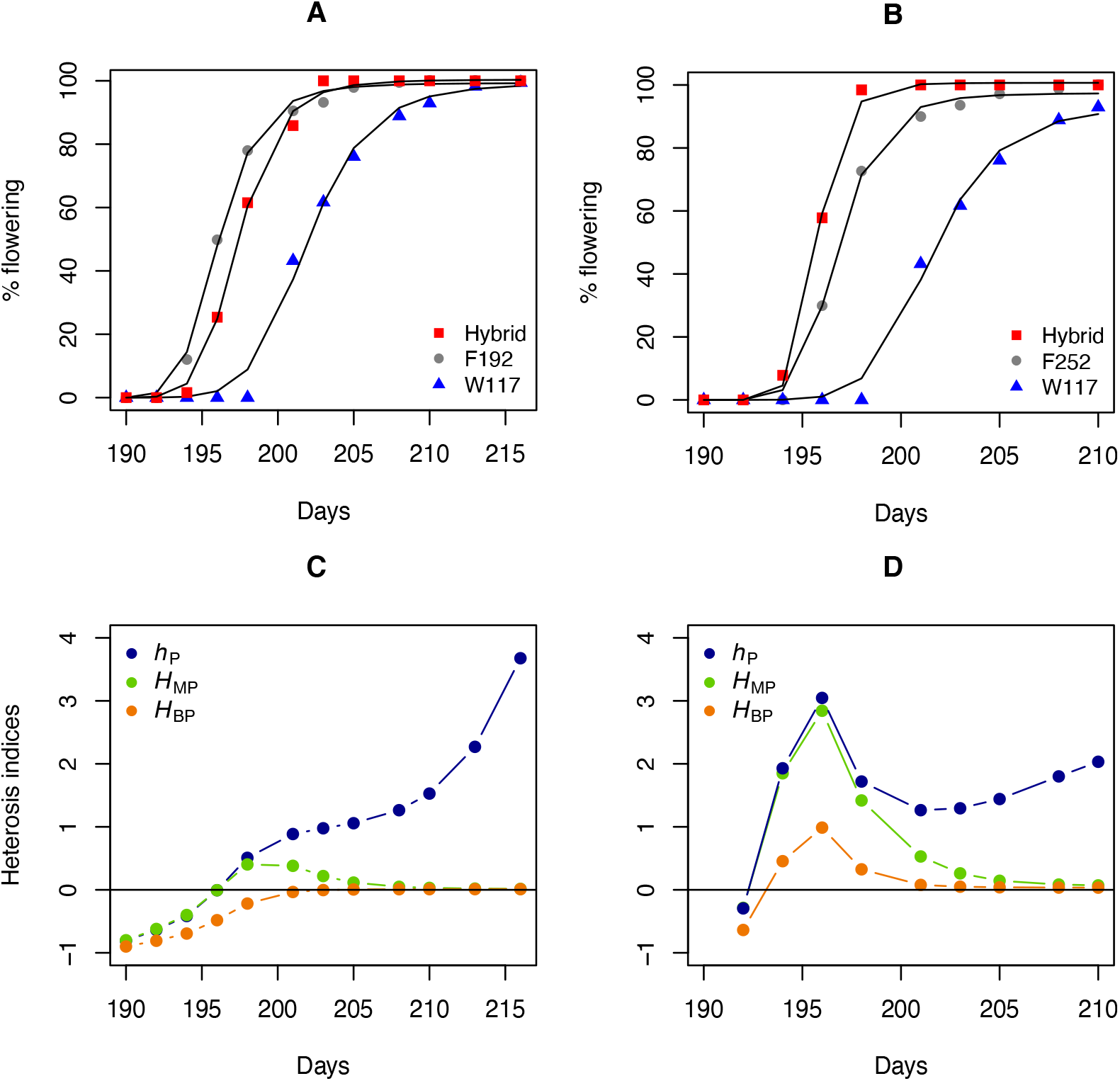
Heterosis for flowering in two maize hybrids. (A) Percentage of flowering over time (number of days since January, 1^st^) for parents W117 and F192 and their hybrid, adjusted with a Hill function. Percentage of flowering over time for parents W117 and F252 and their hybrid. (C) Variations in the heterosis index values for the W117×F192 cross. (D) Variations in the heterosis index values for the W117×F252 cross.

## Discussion

If *H*_MP_ and *H*_BP_ (and their non-normalized forms *H*_mp_ and H_bp_) do not provide reliable information on non-additivity, why are they so commonly used? There are probably both historical and technical reasons: (i) The first scientists who quantified heterosis were plant breeders (Shull, 1908; East, 1936). From an economic perspective, the goal was, and still is, to develop hybrids that are ‘‘better” than the best-or mid-parent values for the desired agronomic traits, and not to know where the hybrid value is relative to the parental values. Heterosis indices have been defined accordingly and the habit has remained; (ii) The indices giving the right non-additivity values, *h*_P_ (= *H*_PR_) for heterosis and *D*_W_ or *D*_F_ for dominance, can take on high to very high values when the parents are close, due to the small differences *z*_2_ – *z*_1_ in the denominator of the fractions. This can produce extreme values that are not easy to represent and manipulate for statistical treatments. Nevertheless, such values are biological realities that convey precisely the inheritance of the traits under study, something that *H*_MP_, *H*_BP_, *H*_mp_ and *H*_bp_ do not. In addition, from a practical point of view, a single index is sufficient to know the position of the hybrid relative to the mid-or best-parent, whereas in a number of studies the authors compute and comment both *H*_MP_ and *H*_BP_ (or *H*_mp_ and *H*_bp_). More importantly, to compare the amplitude of heterosis between traits, developmental stages, crosses or environmental conditions, there is no other choice but to use the only heterosis index – *H*_PR_ – that is not affected by the scale of the parental values and that accounts for the position of the hybrid in the parental range.

## Acknowledgements

We thank our colleagues Dr. Melisande Blein-Nicolas, Dr. Michel Zivy, Dr. Judith Legrand and Pr. Christine Dillmann for their helpful comments on the manuscript, and Dr. Helene Citerne for English corrections. We are grateful to the key persons at the INRA Station of Saint-Martin-de-Hinx, of Euralis and of MASseeds for the 2014 and 2015 field experiments. These experiments were funded by the French Agence Nationale de la Recherche (*Amaizing* project ANR-10-BTBR-01).

## Author contributions

Conceptualization: DdV. Maize experiments: JBF. Data analyses and numerical applications: DdV and JBF. DdV and JBF wrote the manuscript.

## Supplementary data

Heterosis indices: What do they really measure?

Dominique de Vienne and Julie B. Fiévet

**Table S1.** Heterosis indices expressed as functions of genetic effects. Subscripts: same as in Table 1. *μ*, mean of the multilocus homozygous genotypes; 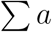, sum of the additive effects; 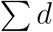, sum of the dominance effects; 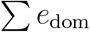, sum of the dominance-by-dominance epistatic effects; 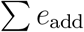, sum of the additive-by-additive epistatic effects; 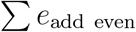, sum of the additive-by-additive epistatic effects involving an even number of genes; 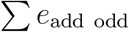, sum of the additive-by-additive epistatic effects involving an odd number of genes (from Fiévet *et al*., 2010).

| Reference | Index | Index as function of genetic effects |
| --- | --- | --- |
| Mid-parent | $H_{\text{mp}} = z_{12} - \bar{z}$ | $\sum d + \sum e_{\text{dom}} - \sum e_{\text{add even}}$ |
| | $H_{\text{MP}} = \frac{z_{12} - \bar{z}}{\bar{z}}$ | $\frac{\sum d + \sum e_{\text{dom}} - \sum e_{\text{add even}}}{\mu + \sum e_{\text{add even}}}$ |
| | $H_{\text{PR}} = \frac{z_{12} - \bar{z}}{(z_2 - z_1)/2}$ | $\frac{\sum d + \sum e_{\text{dom}} - \sum e_{\text{add even}}}{\sum a + \sum e_{\text{add odd}}}$ |
| Best-parent | $H_{\text{bp}} = z_{12} - z_2$ | $\sum d + \sum e_{\text{dom}} - \sum a - \sum e_{\text{add}}$ |
| | $H_{\text{BP}} = \frac{z_{12} - z_2}{z_2}$ | $\frac{\sum d + \sum e_{\text{dom}} - \sum a - \sum e_{\text{add}}}{\mu + \sum a + \sum e_{\text{add}}}$ |

**Fig. S1.**
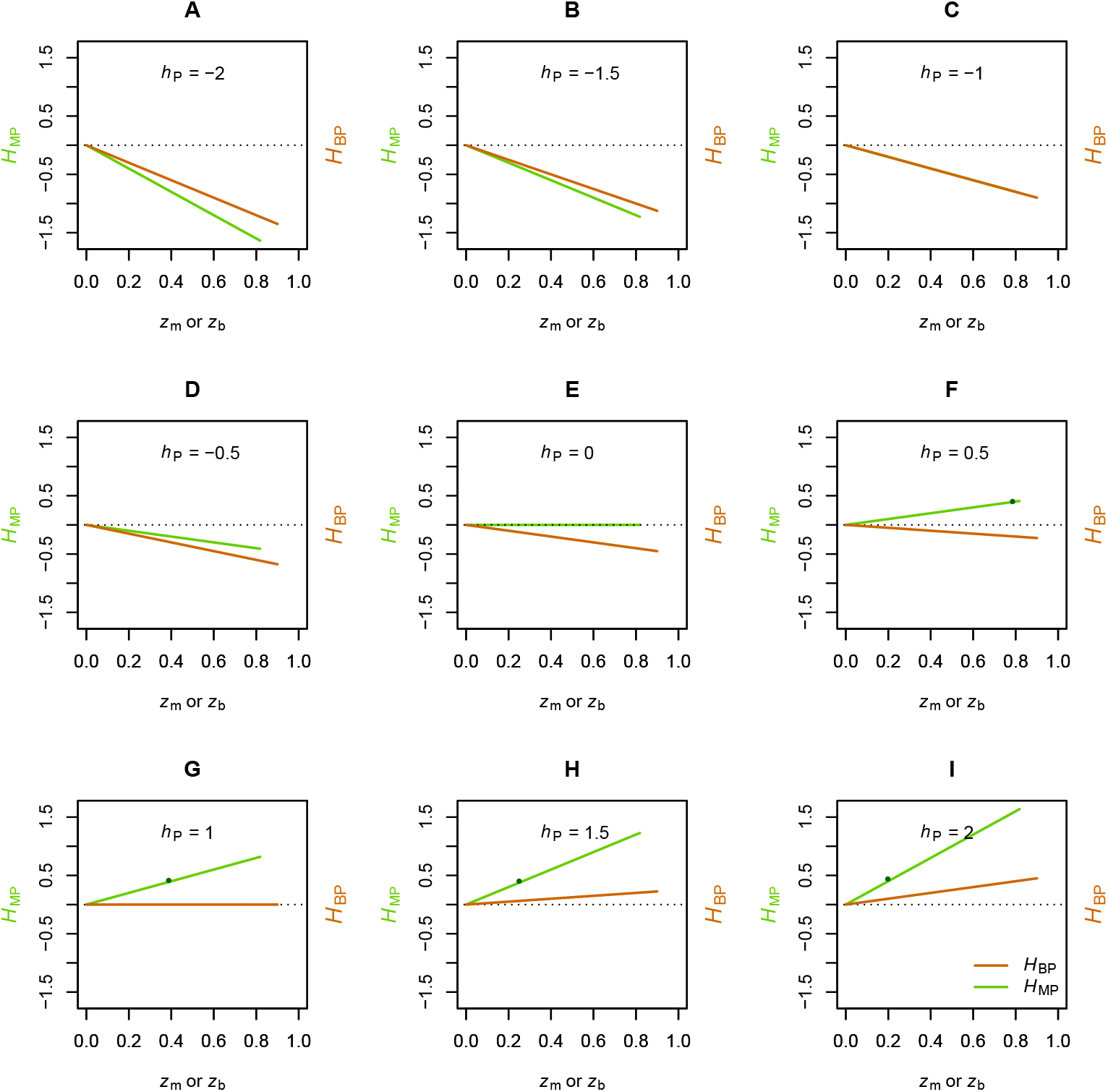
Influence of the scale of the parental values on H_MP_ and H_BP_ for different values of the potence ratio *h*_P_. (A) to (I) *h*_P_ values from –2 to 2. 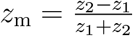 and 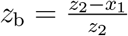, with *z*_1_ = 1 and *z*_2_ varying from 1 to 10 (see equations 1 and 2 in the text). Green line: relationship between *z_m_* and *H*_MP_. Orange line: relationship between *z*_b_ and *H*_BP_. Dotted line: *H*_MP_ or *H*_BP_ = 0. The dark green points show that a given *H*_MP_ value (æ 0.4) can be observed for very different *h*_P_ values, and the same is true for *H*_BP_.

**Fig. S2.**
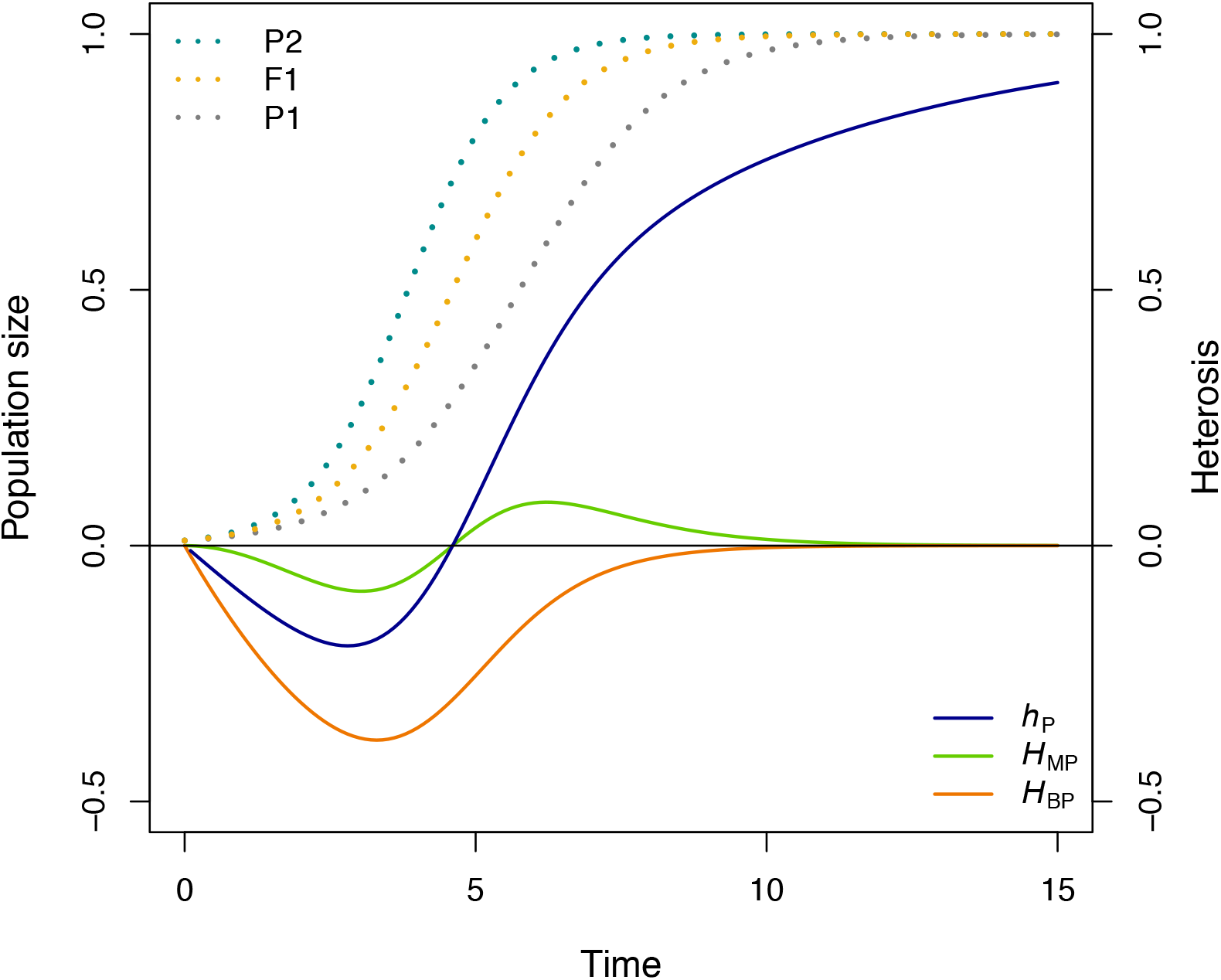
Heterosis for population size (simulations). Population sizes (dotted curves) follow over time a logistic function with *K* =1 and *a* = 100 (see text). Parents P1 and P2 and hybrid F1 have respectively growth rates *r* = 0.8, *r* = 1.2 and *r* = 1 (i.e. this parameter is considered to be additive). Solid curves: profiles of heterosis indices (right scale).

